# Spectral network analysis illuminates coordinated plant traits across a climate gradient

**DOI:** 10.1101/2025.09.18.676927

**Authors:** Rishav Ray, Brandie Quarles-Chidyagwai, Jessica Lyons, Sarah Ashlock, Jennifer R. Gremer, Julin N. Maloof, Troy S. Magney

## Abstract

- Understanding how plant populations respond to environmental variation through functional leaf traits remains challenging due to limitations of traditional phenotyping approaches. Hyperspectral reflectance offers a rapid, non-destructive and high-throughput method to capture functional trait variation and detect signatures of local adaptation across populations.
- We combined hyperspectral data, inverse modeling, and network analysis to investigate population-level variation in *Streptanthus tortuosus*. Using a common garden experiment with four geographically distinct populations, we applied partial least square discriminant analysis (PLS-DA) and ridge regression for population discrimination, inverse PROSPECT modeling to estimate leaf biochemical traits, and canonical correlation analysis to examine trait-climate relationships across historical (1900-1994) and recent (1995-2024) periods. We developed a spectral network approach treating wavelength correlations as biologically meaningful trait networks.
- Populations showed distinct, heritable spectral signatures with high classification accuracy. Significant population differences emerged in anthocyanins, carotenoids, chlorophyll, and water content. Trait-climate correlations shifted between time periods, consistent with historical climate adaptation. Network analysis revealed population-specific integration patterns, with more variable environments displaying greater spectral modularity.
- Hyperspectral signatures provide a high-throughput tool for detecting population-level adaptation and trait coordination. Our findings provide a framework to investigate how plant populations respond to climate change through evolved shifts in trait networks rather than isolated traits alone.

## Introduction

Climate change is driving unprecedented environmental shifts that are reshaping selective pressures on plant populations worldwide (Gienapp *et al*., 2014; Edelsparre *et al*., 2024), yet our ability to detect and understand evolutionary responses to these changes remain limited by traditional approaches of studying adaptation (Franks *et al*., 2014; Anderson & Gezon, 2015; Anderson *et al*., 2025). While common garden experiments and reciprocal transplants have provided foundational insights into local adaptation (Leimu & Fischer, 2008; Hereford, 2009; Schwinning *et al*., 2022), these methods are often constrained by logistical challenges, long generational times, and the difficulty of measuring multiple traits simultaneously across large populations (Merilä & Hendry, 2014; Song *et al*., 2021). The growing availability of genomic data has revolutionized our understanding of genetic basis of adaptation (Savolainen *et al*., 2013; Tigano & Friesen, 2016), but a critical gap remains between high resolution genomic data and the complex, multi dimensional phenotypic data that are targets of natural selection (Houle *et al*., 2010; Furbank & Tester, 2011). This phenotyping bottleneck is particularly acute in the study of climate adaptation, where responses may involve coordinated changes across multiple physiological and biochemical traits rather than simple modifications of individual traits (Nicotra *et al*., 2010; Anderson *et al*., 2012; Hein *et al*., 2021). Consequently, there is an urgent need for high throughput, integrative approaches that can capture the complexity of adaptive phenotypes while scaling to the spatio-temporal dimensions over which climate driven evolution operates (Des Roches *et al*., 2018; Cavender-Bares *et al*., 2022).

Hyperspectral reflectance analysis offers a promising solution to these challenges by providing a rapid, non-destructive window into plant functional traits that integrates information across multiple physiological and biochemical processes (Ustin & Gamon, 2010; Cavender-Bares *et al*., 2016; Wong *et al*., 2023; Wong, 2023; Magney, 2025). Unlike traditional vegetation indices that rely on a few broad spectral bands, hyperspectral data captures reflectance across hundreds to thousands of narrow wavelengths, encoding detailed information about leaf biochemistry, water content, and structural properties (Kokaly *et al*., 2009; Serbin *et al*., 2014). This rich spectral information can be used with models run in inverse mode such as PROSPECT, which enable the quantitative estimation of specific biochemical traits, such as chlorophyll, carotenoids, anthocyanins, leaf-mass per area (LMA), water content, and leaf structure from reflectance spectra (Jacquemoud & Baret, 1990; Féret *et al*., 2021). Recent advances have demonstrated the power of hyperspectral approaches for detecting functional diversity within and among plant communities (Asner *et al*., 2015; Schweiger *et al*., 2018; Wong *et al*., 2023; Li *et al*., 2023; Pau *et al*., 2025), mapping plant stress responses (Gamon *et al*., 2019), linking spectra to gene expression (Chen *et al*., 2025), and even predicting ecosystem function from spectral diversity patterns (Wang *et al*., 2018; Cavender-Bares *et al*., 2022). However, most applications have focused on species-level differences or broad ecological patterns, with limited exploration of how hyperspectral signatures might reveal population-level adaptation and the evolutionary processes that shape functional trait variation within species (Singh *et al*., 2015; Meireles *et al*., 2020). Furthermore, traditional analyses treat spectral bands as independent variables, potentially missing emergent properties that can be gleaned from the coordinated relationship between them (Gamon *et al*., 2020).

Understanding how plants respond to environmental variation requires moving beyond individual traits to consider the coordinated relationships among multiple functional characteristics, a perspective rooted in the concept of phenotypic integration from quantitative genetics and evolutionary developmental biology (Cheverud, 1996; Armbruster *et al*., 2014). Traits rarely evolve in isolation; instead, they are connected through genetic correlations (linkage disequilibrium), developmental constraints, and functional dependencies that result in observable phenotypes (Wagner & Altenberg, 1996; Klingenberg, 2008; He *et al*., 2020; Li & He, 2024). The structure of these trait correlations itself is an evolvable property that can be shaped by natural selection, with different patterns of integration. This can range from highly modular to tightly integrated, representing alternative adaptive strategies for responding to environmental challenges (Hansen & Houle, 2008; Murren, 2012; Wang *et al*., 2023). Network theory has emerged as a powerful framework for understanding such complex systems across biology, from gene regulatory networks that control development (Barabási & Oltvai, 2004; Davidson & Erwin, 2006), to ecological networks that structure community interactions (Bascompte & Jordano, 2007; Olesen *et al*., 2007; Gladstone-Gallagher *et al*., 2023). In each case, network topology, that is the pattern of connections among nodes, provides insights into system function, stability, and evolutionary potential that are undetectable when components are studied in isolation (Albert & Barabási, 2002; Proulx *et al*., 2005). However, applying network approaches to study phenotypic integration has been limited by the challenge of measuring sufficient numbers of traits to characterize network structure, particularly in studies of natural populations where high-throughput phenotyping methods are essential (Murren, 2012; Messier *et al*., 2017; Li & He, 2024). Hyperspectral data offers an unprecedented opportunity to examine trait networks at high resolution and test fundamental hypotheses about how environmental variation shapes the evolution of phenotypic integration (Cavender-Bares *et al*., 2016; Schweiger *et al*., 2018).

The California mountain jewelflower, *Streptanthus tortuosus* (Brassicaceae), provides an ideal system for testing these novel approaches. While most research on local adaptation in *Streptanthus* has focused on other members of the clade, few studies have characterized population-level variation in *S. tortuosus* across California’s diverse climatic gradients (Preston, 1991, 1994; Gremer *et al*., 2020b,a). This species’ wide distribution across elevation and precipitation gradients, combined with common garden experiments, makes it particularly suitable for investigating how environmental variation drives complex trait architecture (Gremer *et al*., 2020a; Bontrager *et al*., 2025). In this study, we combine multivariate hyperspectral reflectance analysis, inverse modeling, and network theory to investigate how population-level variation in spectral traits associates with climate of origin in *S. tortuosus.* Using a common garden experiment spanning four *S. tortuosus* populations, we test whether spectral reflectance data can detect heritable variation in functional traits and reveal population-specific patterns of trait coordination. We apply a novel two-step analytical framework – partial least square discriminant analysis (PLS-DA) for population discrimination followed by ridge regression analysis to identify key wavelength regions – to characterize population specific spectral signatures. Through inverse PROSPECT modeling, we estimate six key leaf functional traits and examine whether their relationships with historical and recent climate variables using canonical correlation analysis across two temporal scales. We then developed a spectral network approach that treats wavelength correlations as biologically meaningful trait networks, revealing distinct patterns of spectral integration among populations. Finally, we test whether these network architectures correlate with long-term climate variability patterns, providing insights into how environmental variation shapes both individual traits and their coordination.

## Materials and Methods

### Plant material and common garden

We conducted a common garden experiment at the UC Davis Vegetable Crops fields in Davis, CA USA, with seeds collected from four populations of *S. tortuosus* (Table 1). Seedlings from these populations were planted at the garden in raised beds with 21 replicates for BH, CC, DPR populations, and 98 replicates for TM2 population. Seeds were stratified for four weeks at 4°C in the dark in a cold room at UC Davis. On October 30, 2023, after stratification, plants were moved to a growth chamber with 12-hour day length and warm light conditions (light intensity ∼100uMol/microeinstein) to stimulate germination. Plants were initially exposed to 22°C day/8°C night temperatures. To harden plants to the cold temperatures of the field, temperatures were adjusted accordingly: 20°C day/7°C night 7-days prior to field transplant, 19°C day/6°C night 6-days prior to field transplant, and 18°C day/5°C night 5-days prior to field transplant. Seedlings were transplanted into a randomized block design with 12 blocks spread across 6 raised beds and 30-cm between experimental plants. The raised beds were 12” tall, 110cm wide, and 14m in length; there was 1m between each bed. Ron’s mix, sand, and perlite were used to fill each bed. The beds were watered with drip irrigation the day before transplant for 5 hours, during the afternoon after planting for 5 hours, and five days after transplant for 5 hours.

**Table 1:** *S. tortuosus* population names and codes used in the study along with their geographic coordinates.

| Population Name | Population Code | Latitude | Longitude |
| --- | --- | --- | --- |
| Table mountain 2 | TM2 | 39.59255 | -121.55072 |
| Ben Hur | BH | 37.40985 | -119.96458 |
| Canyon Creek | CC | 39.58597 | -121.43311 |
| Drum Powerhouse Road | DPR | 39.22846 | -120.81518 |

### Hyperspectral data collection and processing

Leaf-level hyperspectral reflectance measurements were collected from plants grown in the common garden using a full-range spectrometer (HR-1024i; Spectra Vista Corporation—SVC). Spectral reflectance was measured from 400 to 2500 nm with a spectral resolution of 1.5 nm (250–1000 nm), 3.8 nm (1000–1890 nm), and 2.5 nm (1890–2500 nm). Leaf spectra were collected with a leaf clip (LC-RP PRO; SVC) attached to a fiber optic cable. The leaf clip houses an internal tungsten halogen lamp that was standardized for reflectance using a built-in Spectralon white reference panel (Labsphere Inc.). Leaf measurements were carried out on flowering plants with young fully expanded leaves that were big enough to fit the leaf clip; three measurements were taken per plant. This resulted in a total of 153 measurements (BH: 5×3, CC: 7×3, DPR: 7×3, TM2: 32×3). Reference measurements were made before every plant and dark current corrections were automatically taken following each scan with an internal shutter. Spectra were then averaged across the three measurements per plant and smoothed using a Savitzky-Golay filter with a 5-point moving window.

### Statistical analysis

Partial Least Squares Discriminant Analysis (PLS-DA) was performed (*mixOmics* package in R; Rohart *et al*., 2017) to assess population-level differentiation in hyperspectral space. PLS-DA was chosen because it is specifically designed to handle high-dimensional datasets where the number of variables (wavelengths) exceeds the number of samples, a common challenge in hyperspectral analysis. PLS-DA reduces dimensionality while maximizing the separation between groups, making it ideal for identifying population-specific spectral discrimination. It was followed by a PERMANOVA test (*vegan* package; Oksanen *et al*., 2025) to statistically test for differences in spectral data among populations. We used PLS-DA to identify canonical variates that maximally separated the four populations while explaining spectral variance. Broad-sense heritability (H²) was calculated for each wavelength using variance components from a mixed-effects model using the *lme4* package with population as a random effect: H² = σ²G/(σ²G + σ²E), where σ²G is genetic variance among populations and σ²E is environmental/residual variance within populations.

Ridge regression models were fitted for each population(*glmnet* package; Tay *et al*., 2023) to identify wavelength-specific contributions to population classification while avoiding overfitting that commonly occurs with high-dimensional spectral data. The L2 loss function in ridge regression shrinks coefficients toward zero but retains all wavelengths, allowing us to quantify the relative importance of different spectral regions for population classification. Spectral coefficients were extracted across all wavelengths and integrated within six functionally relevant spectral regions (carotenoids: 450-520 nm, chlorophyll: 650-700 nm, NIR plateau: 700-1300 nm, red edge: 680-750 nm, SWIR (short wave infra-red) 1 : 1400-1500 nm, SWIR 2: 1900-2000 nm). To avoid overfitting and provide unbiased performance estimates, we used leave-one-out cross-validation (LOOCV). For each sample, the ridge regression model was trained on all remaining samples, and predictions were made on the held-out sample. Area under the ROC curve (AUC) was calculated from these out-of-fold predictions. Confidence intervals were estimated using 1000 bootstrap replicates of the LOOCV procedure (Fig. S1).

Inverse PROSPECT modeling was applied using the *prospect* R package (Féret & Boissieu, 2024) to estimate six leaf functional traits: anthocyanin content, carotenoid content, chlorophyll content, leaf mass area, structural parameter (N), and equivalent water thickness (estimated water mass per leaf area). Climate data for each population’s origin site were obtained from Flint Basin Characterization Model (Flint *et al*., 2021; Fig. S2), spanning 1900-2024 and divided into historical (1900-1994) and recent (1995-2024) periods. Canonical Correlation Analysis (CCA) was performed (*mixOmics* package) to examine relationships between estimated leaf traits and climate variables across different temporal periods. CCA examines multivariate relationships between multiple leaf traits and multiple climate variables simultaneously, capturing complex trait-environment associations that would be missed by univariate approaches. It identifies linear combinations of traits and climate variables that are maximally correlated, revealing coordinated trait response to environmental gradients.

To address spectral autocorrelation and reduce redundancy for the network analysis, we implemented a filtering strategy based on spectral information content. We calculated the Shannon diversity index (H = -Σ(pi × ln(pi))) for each spectrum to estimate the effective number of independent wavelengths, where pi represents the proportion of total reflectance at wavelength i (Fig. S3). The mean Shannon index across all samples was 7.3, yielding an effective number of wavelengths (eH) of approximately 1495. Combined with our heritability filtering (H² > 0.5), this approach retained 987 wavelengths with both genetic differentiation and non-redundant spectral information, acknowledging that adjacent wavelengths are inherently correlated due to the physical properties of light-matter interactions. This filtering strategy reduced computational burden while preserving biologically meaningful spectral variation for network construction. For each population, correlation matrices were calculated between all wavelength pairs using Pearson correlation coefficients. To account for sample size limitations, we employed a bootstrap approach, randomly sampling 10 individuals with replacement 100 times and constructing networks for each iteration. Edges were retained only for correlations exceeding |r| > 0.7. Four key network parameters were calculated using the *igraph* package: (1) Density -the proportion of possible edges that are actually present in the network, calculated as the number of observed edges divided by the maximum possible number of edges (n(n-1)/2 for undirected networks). Values range from 0 (no connections) to 1 (fully connected), with higher values indicating more integrated spectral responses across wavelengths. (2) Centralization -a measure of how unequally distributed the connections are across nodes, quantifying the extent to which the network is dominated by highly connected “hub” wavelengths. Centralization is calculated based on the variance in node degrees relative to a star network (maximum centralization) and ranges from 0 (all nodes equally connected) to 1 (one central hub connected to all other nodes). High centralization indicates that spectral integration is controlled by a few key wavelengths. (3) Modularity -a measure of community structure that quantifies how well the network can be divided into distinct groups of densely connected nodes (modules) with sparse connections between groups. Modularity was calculated using the Louvain algorithm, which optimizes the modularity function Q = Σ[Aij - (kikj/2m)]δ(ci,cj), where Aij is the adjacency matrix, ki is the degree of node i, m is the total number of edges, and δ(ci,cj) = 1 if nodes i and j belong to the same community. Values range from -0.5 to 1, with higher values indicating stronger modular organization of spectral responses. (4) Transitivity -clustering coefficient -the probability that two nodes connected to a common neighbor are also connected to each other, measuring the tendency for wavelengths to form tightly connected clusters. Calculated as the ratio of observed triangles to possible triangles in the network: C = 3 × (number of triangles) / (number of connected triples). Values range from 0 to 1, with higher values indicating greater local clustering of spectral correlations. CCA was performed between population-level network metrics and long-term climate variability measures (coefficients of variation for temperature, precipitation, and climatic water deficit from 1900-2024) to test relationships between network topology and environmental variation.

All analyses were performed in R (version 4.2.1; R Core Team, 2022). Significance testing for network metrics was conducted using ANOVA with Tukey’s HSD post-hoc tests. Bootstrap confidence intervals were calculated for all network parameters. Statistical significance was set at α = 0.05, with Bonferroni correction applied for multiple comparisons where appropriate.

## Results

### Population-level hyperspectral differentiation and genetic variation

To determine whether *S. tortuosus* populations maintain distinct spectral signatures under standardized common garden conditions, we performed PLS-DA on the hyperspectral data over a multivariate space. The PLS-DA ordination demonstrated significant differences in spectral values among populations (PERMANOVA F:17.43, p-value:0.001), with the first two canonical variates explaining 97.93% of the total spectral variation (Fig. 1A, S4). The populations formed distinct clusters in the ordination space, with TM2 and DPR populations showing greatest separation along the first canonical variate, while the BH population separated in a distinct cluster along the second canonical variate consistent with its geographical separation (Fig. 1B).

**Fig. 1:**
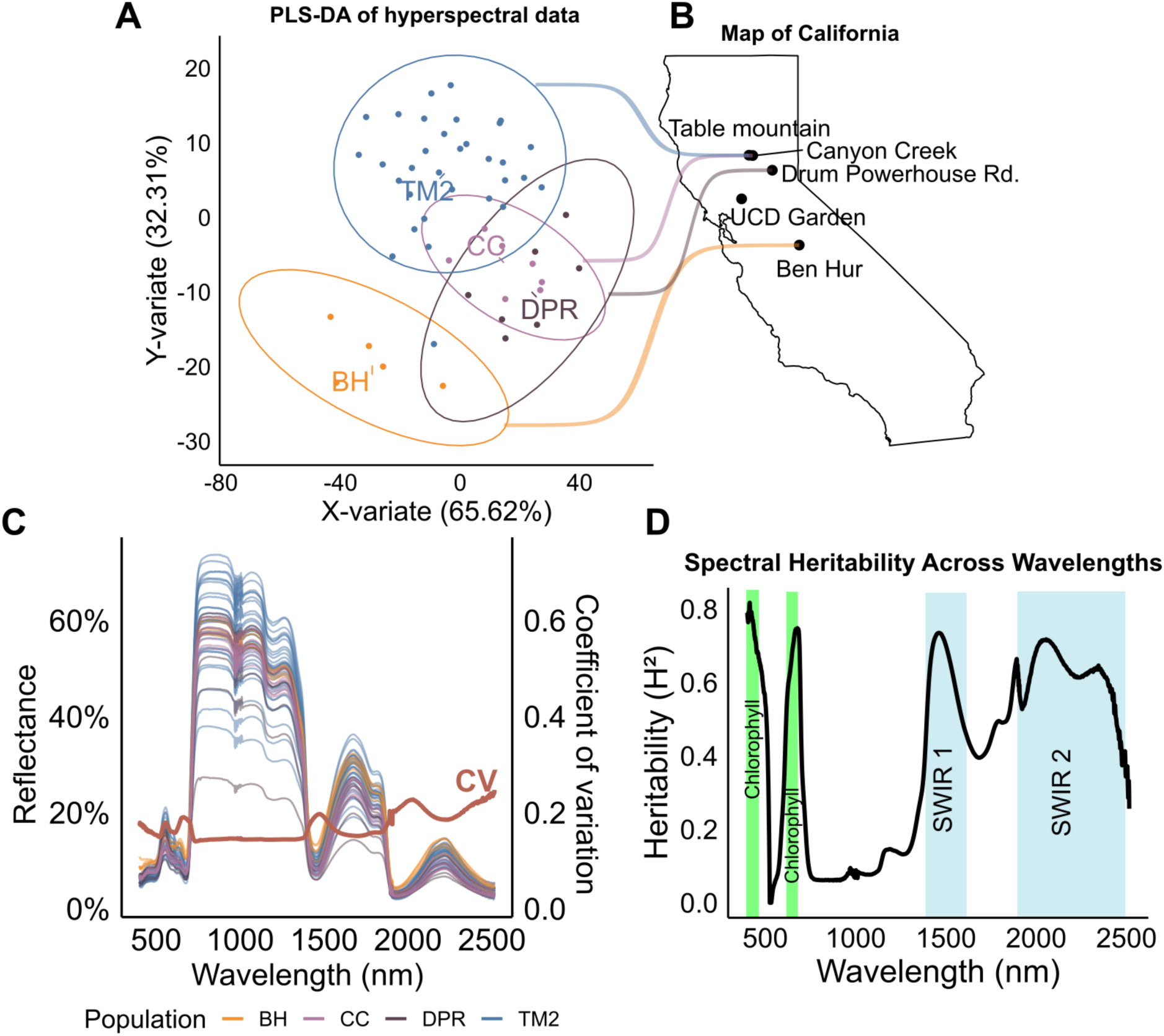
Hyperspectral variation and heritability in *Streptanthus tortuosus* populations across California. **A)** Partial Least Squares Discriminant Analysis (PLS-DA) of hyperspectral data showing four *S. tortuosus populations* (TM2, CC, DPR, and BH) along two principal variates. The x-variate explains 65.62% of the spectral variation while the y-variate explains 32.31%. **B)** Map of California showing the geographic distribution of the sampled populations: Table Mountain (TM2), Canyon Creek (CC), Drum Powerhouse Road (DPR), Ben Hur (BH) and the UC Davis Common Garden (UCD) where all these populations were planted and measured. **C)** Reflectance spectra by population across wavelengths from 400-2500 nm. Different coloured lines represent individual plant measurements. The red line (labeled “cv”) shows the coefficient of variation across wavelengths (right y-axis). **D)** Broad-sense heritability (H²) estimates across wavelengths from 400-2500 nm, showing variation in the genetic contribution to spectral reflectance traits. The coloured bands indicate spectral regions that correlate with leaf chlorophyll content (green), and SWIR regions (blue).

Despite being measured under the same common garden conditions, the samples showed spectral variation across key regions of the wavelength range, denoted by the coefficient of variation (CV) plotted in the second axis (Fig. 1C). The CV across samples peaked in several key spectral regions, with the highest variability observed around 680 nm (chlorophyll absorption), and water absorption bands around 1400 nm and 1900 nm; regions that correspond to functionally important leaf properties. Broad-sense heritability (H^2^), which estimates the amount of phenotypic variation explained by genotype, revealed substantial genetic control over spectral reflectance traits (Fig 1D). The H^2^ values peak considerably across the chlorophyll and SWIR regions, suggesting that the genetic variation is concentrated in spectral regions that reflect key physiological and biochemical leaf properties. Taken together, maintenance of population-specific spectral signatures under common garden conditions, combined with high heritability estimates, provides strong evidence for genetic differentiation in leaf optical properties among *S. tortuosus* populations.

### Ridge regression analysis reveals population-specific spectral signatures

Having established that there are heritable differences among populations, we next asked, what are the specific wavelengths that distinguish between the populations? To answer this, we applied ridge regression analysis and quantified the predictive power of hyperspectral data for population classification. Regression coefficients across the spectra (referred to as spectral coefficients hereafter) varied markedly across wavelengths and among populations, with each population exhibiting unique patterns of spectral importance (Fig. 2A). All populations showed pronounced spectral coefficients in the visible region (400-700nm), particularly in areas corresponding to carotenoid (450-520 nm), and chlorophyll (400-250 & 650-700nm) absorption bands. Notable population specific differences emerged in the water absorption regions (1400-1500 and 1900-2000 nm), where DPR displayed stronger spectral coefficients compared to other populations, which varied in those regions (Fig 2C). The near-infrared region (700-1300 nm) is influenced by leaf internal structure and showed distinct coefficient patterns, with BH exhibiting particularly strong signals in this spectral region (Fig. 2A). We numerically integrated the spectral coefficients across six functionally relevant wavelength regions and ranked them to quantify the population specific spectral “fingerprints” (Fig. 2C). TM2 consistently ranked highest in three key regions: chlorophyll, carotenoids, and SWIR 2 absorption bands. In contrast, BH was most sensitive to the near infra-red (NIR) region. CC and DPR populations showed more moderate rankings across most spectral regions, with CC ranking highest in the SWIR 1 region and DPR showing intermediate values across all spectral domains. Furthermore, the discriminative power of hyperspectral data for population identification was high (AUC 0.838-0.99) across all populations, indicating that hyperspectral reflectance data contain sufficient population-specific information to achieve high classification accuracy (Fig 2B, S1). In summary, these results suggest clear population-specific differentiation in spectral signatures, indicating that populations vary in multiple optical properties.

**Fig. 2:**
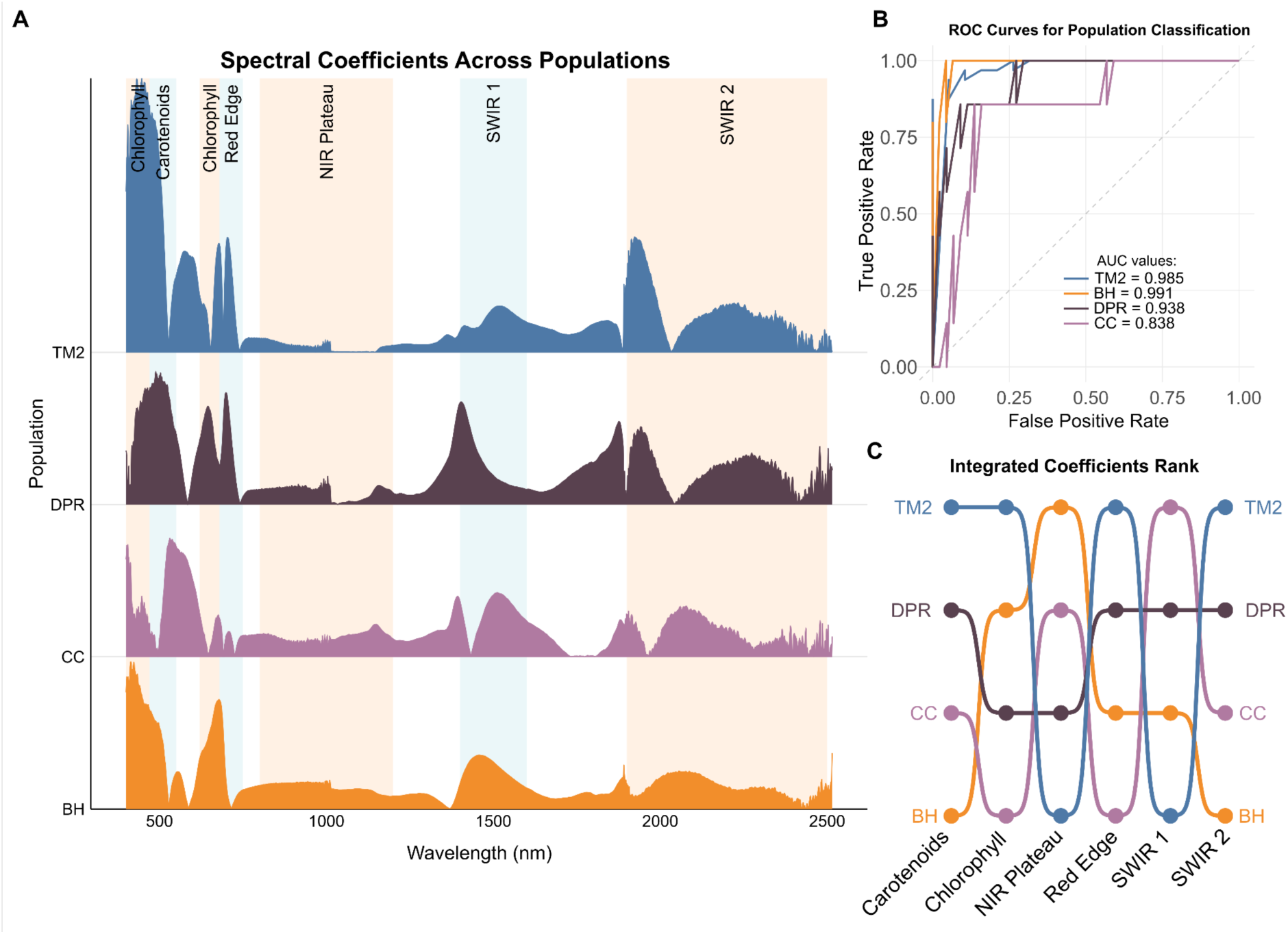
Spectral characteristics and classification performance across *Streptanthus tortuosus* populations. **A)** Spectral coefficients derived from ridge regression across wavelengths (400-2500 nm) for four *S. tortuosus* populations (TM2, DPR, CC, and BH). Background shading highlights key spectral regions associated with different leaf properties: carotenoids, chlorophyll, NIR plateau, red edge, SWIR 1, and SWIR 2. **B)** Receiver Operating Characteristic (ROC) curves showing classification performance for each population based on ridge regression models. All populations demonstrate excellent classification accuracy with Area Under the Curve (AUC) (Fig. S1) values ranging from 0.838 (CC) to 0.99 (BH), indicating high discriminative power of spectral data for population identification. **C)** Integrated coefficients rank plot showing the relative importance of six key spectral regions in distinguishing between populations. Each colored line represents a population (TM2, DPR, CC, and BH), with vertical positions indicating their rank (higher = stronger signal) for each spectral region.

### Leaf trait differences and climate associations across temporal scales

We next asked whether the variation in reflectance in different populations can be explained by various home climatic variables that they have experienced. To that end, we applied the inverse PROSPECT (Jacquemoud & Baret, 1990; Féret & Boissieu, 2024) model to estimate six key leaf functional traits from the hyperspectral data, namely: anthocyanins, carotenoids, chlorophyll, leaf mass area, N (leaf structural complexity), and equivalent water thickness (EWT) (Fig 3A). The traits show significant population-level variation, e.g. anthocyanin contents were highest in BH and CC populations, exceeding levels in DPR and TM2 populations by about two-fold. Carotenoid content showed the opposite pattern, with TM2 displaying markedly higher concentrations compared to other populations, and not detectable from BH spectra. Chlorophyll content was highest in CC, while BH showed substantially lower levels. EWT also varied significantly among populations, with CC again showing the highest values compared to BH. Leaf mass area and the structural parameter N showed more modest but consistent population differences, indicating coordinated variation in both biochemical and structural leaf properties.

**Fig. 3:**
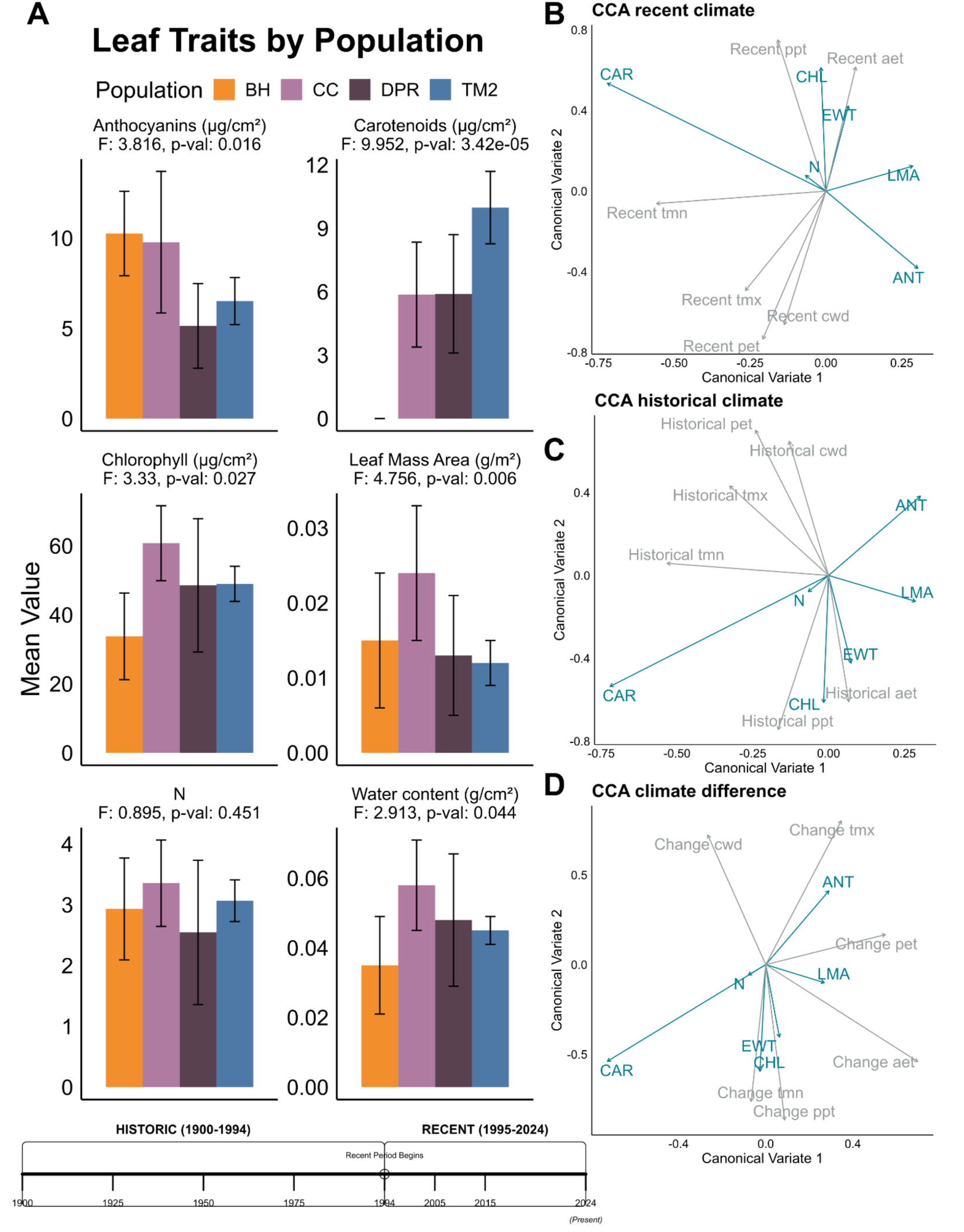
Leaf trait variation across populations and relationships with climate variables. **A)** Mean values (±SE) and F-statistic and p-value of ANOVA test for six key leaf traits across four *S. tortuosus populations* (BH, CC, DPR, and TM2). Traits include anthocyanins (μg/cm²), carotenoids (μg/cm²), chlorophyll (μg/cm²), leaf mass area (g/m²), leaf structure constant (N), and water content (g/cm²). The Timeline at the bottom indicates the historical (1900-1994) and recent (1995-2024) climate periods used in the analysis. **B-D)** Canonical Correlation Analysis (CCA) biplots showing relationships between leaf traits (blue vectors) and climate variables (gray vectors) during recent (**B**) and historical (**C**) time periods, and the difference between the two time periods (**D**). The first canonical variate explains the primary correlation structure between leaf traits and climate conditions.

To test for environmental associations with these estimated traits, we applied canonical correlation analysis with populations’ home climatic variables going back to 1900, split between recent (1995-2024), and historic (1900-1994) time periods. The analysis revealed distinct patterns of trait-climate associations across different temporal periods. For recent climate conditions, the first canonical correlation explained strong relationships between leaf traits and contemporary environmental variables (Fig. 3B). Chlorophyll content and EWT correlated positively with recent actual evapotranspiration and precipitation and negatively with climate water deficit. Historical climate relationships showed similar trait-environment associations compared to recent patterns (Fig. 3C). However, climate change analysis (difference between recent and historical periods) provided insights into how shifting environmental conditions correlate with current trait variation (Fig. 3D). Anthocyanins showed a strong positive correlation with change in maximum temperature, and interestingly, chlorophyll, and EWT both correlated positively with changes in precipitation and minimum temperature patterns. These temporal patterns reveal that trait-climate correlations differ between historical and recent periods, with the strength and direction of associations varying across time intervals.

### Spectral network architecture reveals population-specific network patterns linked to climate variability

We then asked whether the leaf reflectance patterns are co-ordinated amongst themselves and if the relationship between them was population specific or not. To that end, we applied a network approach to build a spectral graph, where each node represents a particular wavelength, and each connecting edge is the correlation coefficient of their reflectance value between the two nodes. This network analysis revealed distinct population-specific network topology with quantitatively significant differences in spectral integration patterns (Fig. 4 A-B). TM2 and DPR populations exhibited dense, highly interconnected networks with numerous edges connecting across wavelengths, reflected in significantly higher network density and degree compared to BH and CC (p < 0.0001). In contrast, BH and CC populations displayed more modular network structures characterized by distinct clusters of highly connected wavelengths separated by sparser connections, with modularity values significantly higher than TM2 and DPR, confirming compartmentalized spectral organization. CC exhibited the highest centralization values, suggesting that spectral integration in this population is dominated by a few highly connected wavelength “hubs.” Transitivity showed more modest but significant differences, with DPR displaying the highest values, indicating greater local clustering of wavelength correlations. These results demonstrate that populations have fundamentally different approaches to organizing spectral trait correlations, ranging from highly integrated (TM2, DPR) to highly modular (CC, BH) network architectures suggesting population-specific variation in the coordination of leaf optical properties.

**Fig. 4:**
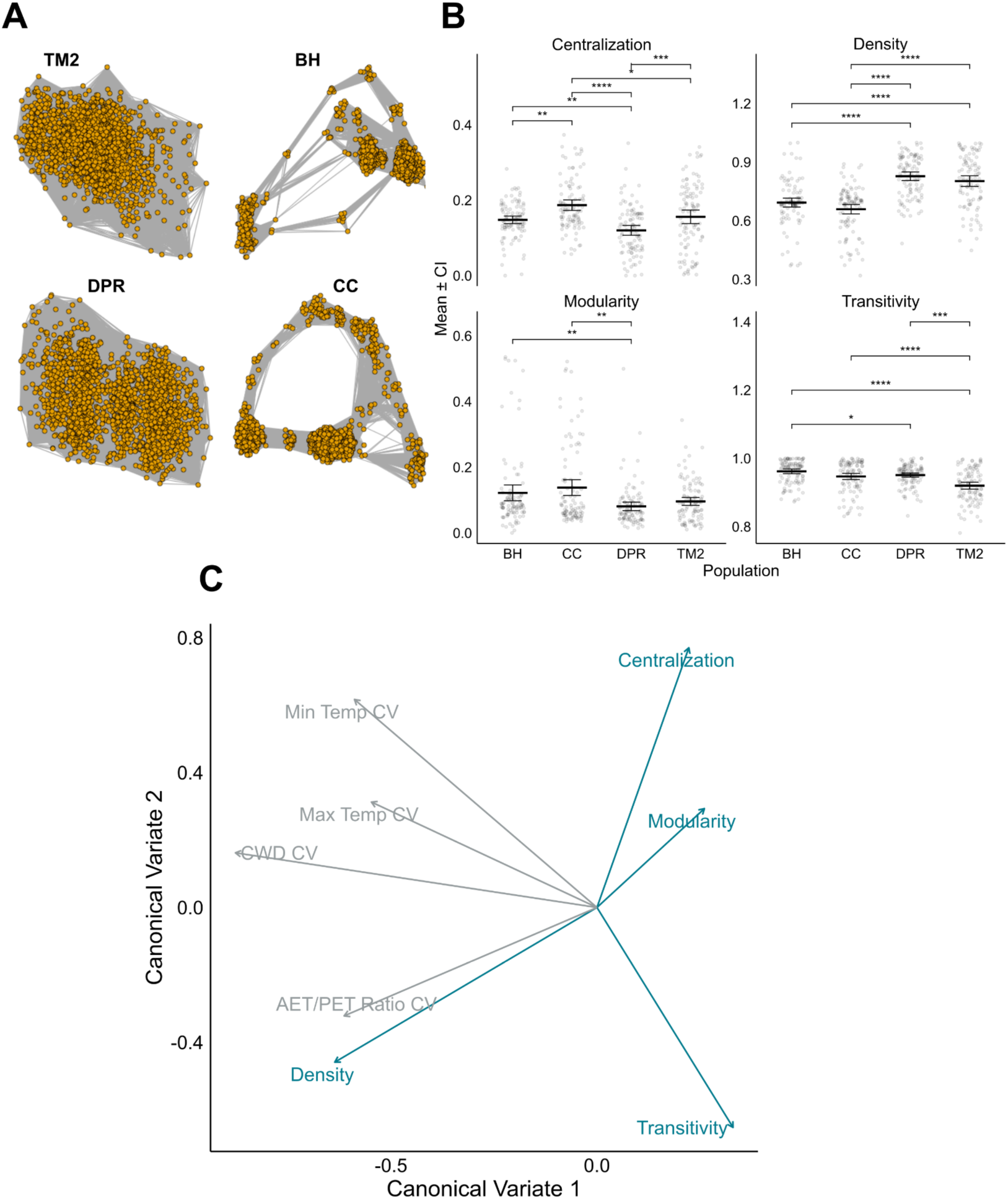
Spectral network analysis and climate-network relationships in *S. tortuosus* populations. **A)** Network visualization of spectral relationships in four *S. tortuosus* populations (TM2, BH, DPR, and CC). Each node represents a wavelength, and edges represent correlations between wavelengths. **B)** Comparison of four network parameters (centralization, density, modularity, and transitivity) across populations. Box plots show the distribution of bootstrapped values (n=100, 10 individuals with replacement per iteration) with significance indicators for population differences (*p<0.05, **p<0.01, ***p<0.001, ****p<0.0001). **C)** CCA biplot showing relationships between network metrics (blue vectors) and climate variability parameters (gray vectors) from 1900-2024.

Canonical correlation analysis revealed that these population differences in spectral network organization are systematically related to long-term climate variability patterns, rather than conditions in specific historical or recent periods (Fig. 4C). The first canonical variate separated density from the other network parameters, while the second canonical variate distinguished centralization and modularity and transitivity. The AET/PET ratio correlated highly with density, suggesting that populations from environments with higher water availability (higher AET/PET) tend to have denser, more integrated spectral networks. Conversely, centralization and modularity both loaded positively on the second canonical variate and were associated with minimum and maximum temperature variability, indicating that populations from thermally variable environments have more centralized and modular spectral integration patterns. Furthermore, The coefficient of variation in climatic water deficit (CWD CV) loaded negatively on the first canonical variate, suggesting that populations experiencing greater water stress variability are associated with lower transitivity and higher density in their spectral networks.

## Discussion

This study demonstrates the power of integrating advanced hyperspectral analysis with network theory to reveal patterns of plant trait coordination. Our novel analytical framework discriminated four *S. tortuosus* populations with good accuracy (AUC ranging from 0.838 -0.99), while revealing that spectral traits represent heritable characteristics. The combination of PLS-DA for population discrimination and ridge regression for identifying key wavelength regions proved highly effective, with population specific signatures observed even under common garden conditions. Beyond individual trait variation, our network analysis uncovered distinct patterns of spectral coordination among populations, ranging from highly connected (TM2) to modular (CC, BH) architectures that correlate with long-term climate variability. Critically, these methodological advances revealed that populations have evolved not only different leaf biochemical properties, with notable differences in anthocyanins, carotenoids, and chlorophyll content, but also fundamentally different strategies for coordinating these traits reflected through their spectral networks. Strong correlations between network topology and historical climate patterns suggest that environmental variability has shaped both individual traits and their patterns of integration, representing a new perspective on the evolution of complex trait architecture in natural populations.

Building on these climate network relationships, specific spectral regions provide additional insights into the functional basis of trait variation. The near-infrared region (700-1300 nm) showed distinct coefficient patterns, with BH exhibiting particularly strong signals in this structurally-sensitive spectral region. The NIR plateau is primarily influenced by leaf internal structure rather than biochemical composition, with reflectance determined by the scattering of light at air-cell wall interfaces within the leaf mesophyll (Jacquemoud & Baret, 1990; Féret et al., 2021). Higher NIR reflectance typically indicates greater leaf thickness, increased intercellular air spaces, or altered mesophyll organization—structural characteristics that can influence light capture efficiency, gas exchange, and water use efficiency (Onoda et al., 2017). The strong NIR signal in BH may reflect population-level variation in structural traits that could be related to environmental conditions at this population’s origin site (Niinemets, 2001; Wright *et al*., 2004; Karabourniotis *et al*., 2021). However, it is important to emphasize that our study cannot distinguish between spectral variation arising from adaptive response to local environment versus genetic drift, founder effects, or population structure. While the functional significance of the estimated traits correlating with the environmental variables suggests that they may be ecologically relevant, we make no causal claims about the evolutionary mechanisms that drive these patterns.

Our hyperspectral network analysis represents a shift from traditional approaches that treat spectral bands as independent variables to a systems-level framework that captures the coordinated relationship between wavelengths. While previous studies have relied primarily on vegetation indices or individual wavelength analysis (Bannari *et al*., 1995; Thenkabail *et al*., 2000; Gamon *et al*., 2019), our network approach reveals emergent properties of spectral trait integration that are not captured by conventional methods. Our two-step analytical pipeline, combining PLS-DA for population discrimination with ridge regression for mechanistic insight, offers significant advantages over single method approaches by first establishing the existence of population differences and then identifying the specific wavelength regions driving these patterns. This framework addresses a key gap in evolutionary ecology, where high throughput phenotyping methods are needed to match the resolution of genomic tools (Houle *et al*., 2010; Furbank & Tester, 2011; Pieruschka & Poorter, 2012; Cavender-Bares *et al*., 2022).The integration of canonical correlation analysis across multiple temporal scales (historic vs. recent climate) further demonstrates how advanced statistical methods can detect trait correlation with environmental variables. This study provides a template for studying evolutionary responses to environmental change, which can be further bolstered by addition of genomic and transcriptomic data ,to study adaptation in real-time (Hoffmann & Sgrò, 2011; Anderson *et al*., 2012; Franks *et al*., 2014; Merilä & Hendry, 2014).

Additionally, our spectral network analysis reveals a previously unrecognized level of biological organization, i.e. the coordinated structure of the reflectance patterns, that parallels network organization observed across diverse biological systems. While reflectance patterns themselves are not direct targets of selection, they represent emergent properties of functionally important traits such as pigment composition, water retention capacity, and leaf structural characteristics that can be under direct selective pressure. Similar to how gene regulatory networks exhibit modular architectures that facilitate evolvability while maintaining stability (Wagner *et al*., 2007; Espinosa-Soto & Wagner, 2010), our spectral networks show population-specific patterns of modularity and integration that are consistent with different strategies for coordinating underlying physiological and biochemical processes. The dense, highly connected networks in TM2 and DPR populations mirror the “small-world” properties observed in metabolic networks, where high clustering facilitates local optimization while maintaining global connectivity (Jeong *et al*., 2000; Barabási & Oltvai, 2004). Conversely, the modular architectures in CC and BH populations resemble the compartmentalized organization of ecological food webs, where modularity enhances stability against perturbations while allowing independent evolution of subsystems (Olesen *et al*., 2007; Stouffer & Bascompte, 2011). This biological network perspective suggests that natural selection acts not only on individual functional traits but also on the patterns of coordination among these traits, with spectral networks providing a high-dimensional window into this complex trait architecture, as proposed in quantitative genetics theory (Cheverud, 1996; Armbruster *et al*., 2014). The correlation between network topology and climate variability patterns supports the hypothesis that environmental unpredictability favors modular organization, thereby allowing rapid reconfiguration of trait combinations, while stable environments favor integrated networks that optimize coordinated responses (Schlosser & Wagner, 2004). Our findings thus extend network thinking beyond its traditional domains in molecular biology and ecology to encompass the evolution of complex phenotypes in the spectral sensing domain. This network perspective offers practical applications for conservation biology, as species with highly integrated spectral networks (low modularity, high density) may be more vulnerable to environmental disruption due to their tightly coupled trait systems.

While our methodological framework demonstrates significant promise, several limitations warrant consideration. First, our analysis was restricted to a single growing season under common garden conditions, which does not fully capture the dynamic of spectral trait expression across environments or temporal variation. This is highlighted by the absence of detectable carotenoid levels in the BH population using the inverse modeling. This highlights potential limitations in our inverse PROSPECT modeling approach for certain biochemical compounds under specific conditions (Féret & Boissieu, 2024). This could reflect either genuine biological differences in carotenoid accumulation strategies or methodological constraints in detecting low-concentration pigments in certain leaf types (Wong, 2023). Additionally, while our network analysis reveals compelling patterns of spectral integration, the biological mechanisms underlying these coordination patterns remain unclear. Future studies combining transcriptomic and metabolomic approaches with hyperspectral analysis could provide mechanistic insights into how plants achieve different integration strategies.

The successful implementation of this analysis framework requires careful consideration of several methodological parameters and quality control practices that can significantly influence results. Key analytical choices include spectral smoothing window size (we used Savitzky-Golay filtering with a 5-point window), correlation threshold for edge retention (|r| > 0.7), and number of bootstrap replicates (n = 100). Each of these parameters represent a trade-off between noise reduction, information retention, and computational complexity that should be optimized for specific dataset and research questions. The small number of populations and relatively small sample size used in this study also limits our ability to generalize across the species’ full geographic range. Future studies with a larger sample size and more populations would significantly strengthen our conclusions about climate trait relationships. Field measurement quality control is another essential component and should include consistent leaf positioning, regular white reference calibration, and systematic checking for spectral artifacts or outliers that could propagate through network analyses (Burnett *et al*., 2021). Importantly, network metrics (density, modularity, centralization, transitivity) and biochemical trait estimates should be interpreted comparatively between treatments or populations rather than as absolute biological values, as their biological meaning emerges from relative patterns rather than specific numerical thresholds. This analytical framework shows considerable promise for scaling to larger spatial and temporal dimensions: canopy-level hyperspectral imaging could extend population comparisons to landscape scales (Kaur *et al*., 2024; Pierrat *et al*., 2025)(, while integration with emerging airborne (Cawse-Nicholson *et al*., 2025) and satellite hyperspectral sensors (Cawse-Nicholson *et al*., 2025; Caplan & Huemmrich, 2025) could enable real-time monitoring of trait network evolution across entire species ranges.

## Conclusion

This study demonstrates how hyperspectral analysis can be integrated with network theory to understand population-level trait coordination in plants. Our analytical framework reveals that spectral data encode information about both individual functional traits and their patterns of coordination, providing a high-throughput approach to studying complex phenotypes in natural populations. Future integration of this spectral framework with genomic and transcriptomic datasets could illuminate the molecular mechanisms underlying different coordination strategies, potentially identifying the genetic basis of trait integration and informing predictive models of population responses to environmental change. Such multi-scale applications will be essential for predicting and monitoring how natural populations respond to accelerating environmental change.

## Acknowledgements

This research was funded by the National Science Foundation (DEB-2129589 to J.N.M, J.R.G, T.S.M, and Denneal Jamison-McClung) and The National Institute of Food and Agriculture, United States Department of Agriculture (USDA-NIFA) award CA-D-PLB-2795-H to JNM. We acknowledge the contribution from Maya Arakaki, technician Kate Ruiz-Cox, and Carlos Perez and all the undergraduate students involved in setting up the field and data collection. J. Schmitt provided valuable guidance on study design, analyses, and manuscript drafts.

## Author contribution

Conceptualization (research direction and analytical framework): RR, JNM, TSM, (common garden and data collection): BQ-C, JRG, JNM, TSM; Data curation: RR, JL; Formal Analysis: RR, JNM, TSM; Funding acquisition: JRG, JNM, TSM; Investigation: RR, JNM, TSM; Visualization: RR; Writing – original draft: RR; Writing – review & editing: RR, BQ-C, JL, SA, JRG, JNM, TSM

## Data availability

Raw data and source code are available in the following Github repository. https://github.com/rishavray/spectral-network

## Supplementary Figures

**Fig. S1:**
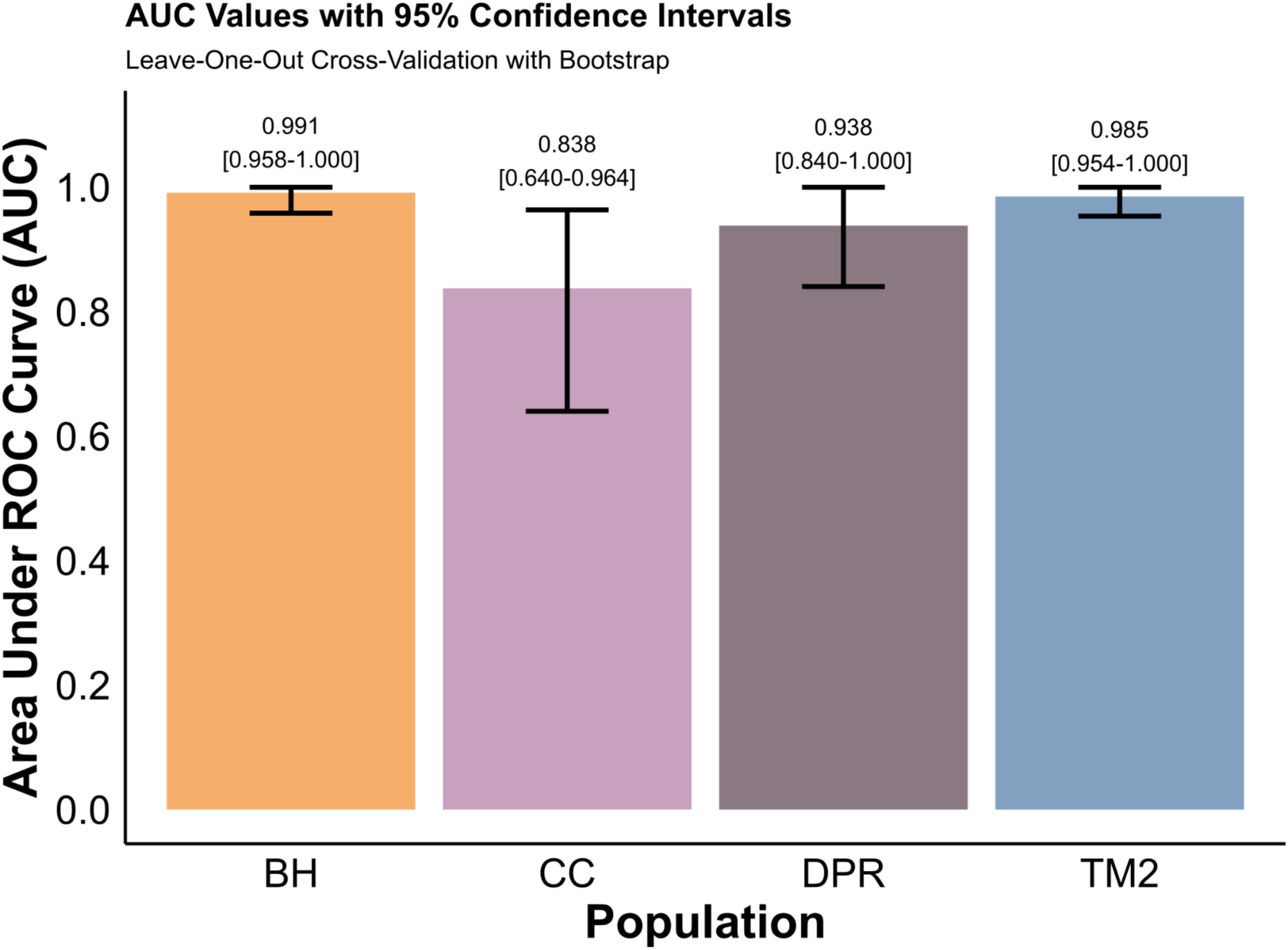
Area under the curve values for each population with 95% confidence intervals calculated with 1000 bootstrap iterations. Bootstrap confidence intervals for AUC values were narrow across all populations, indicating stable and reliable classification performance despite modest sample sizes.

**Fig. S2:**
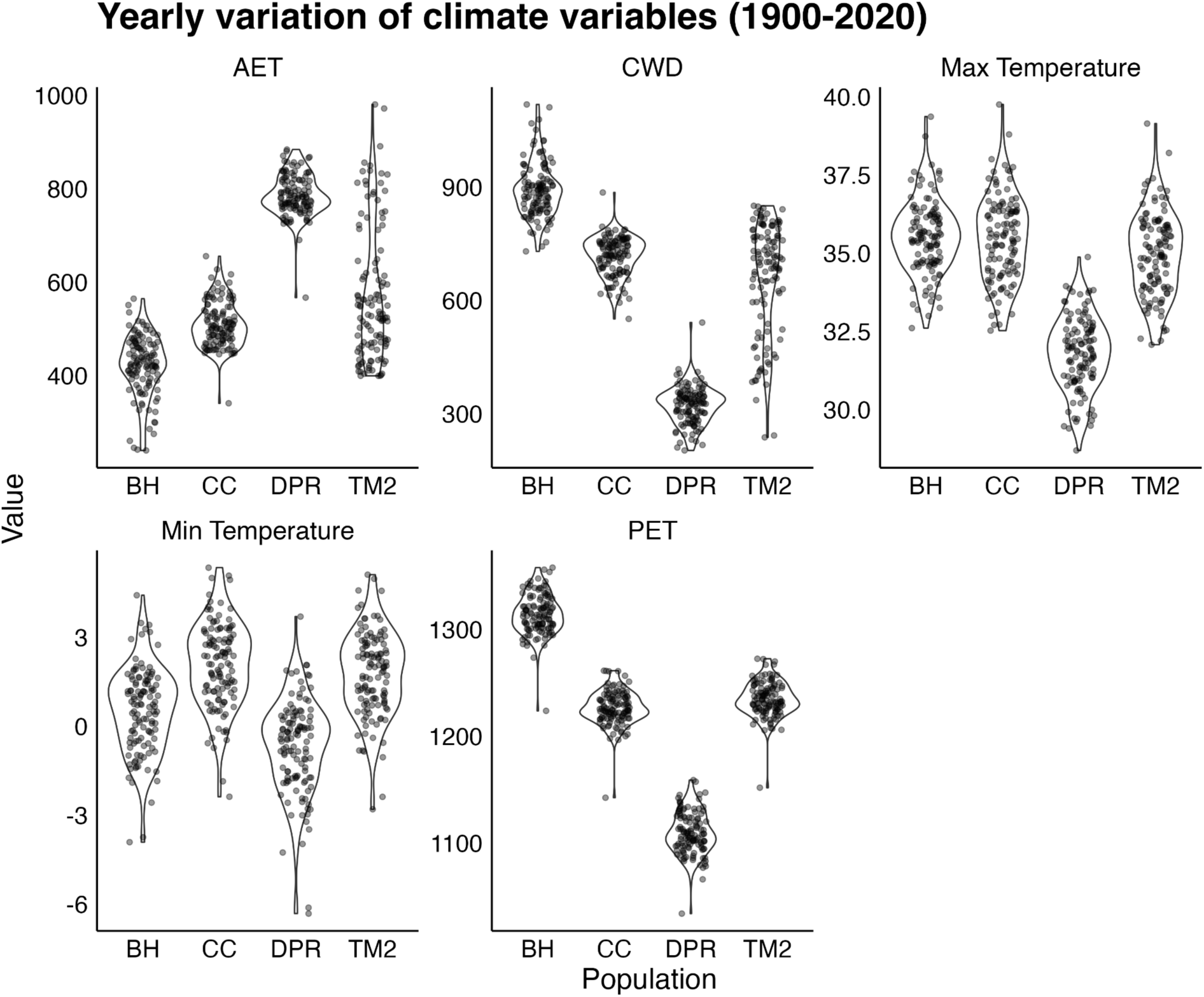
Yearly variation of the five climate variables used in the study. Each dot represents a year for each individual population.

**Fig. S3:**
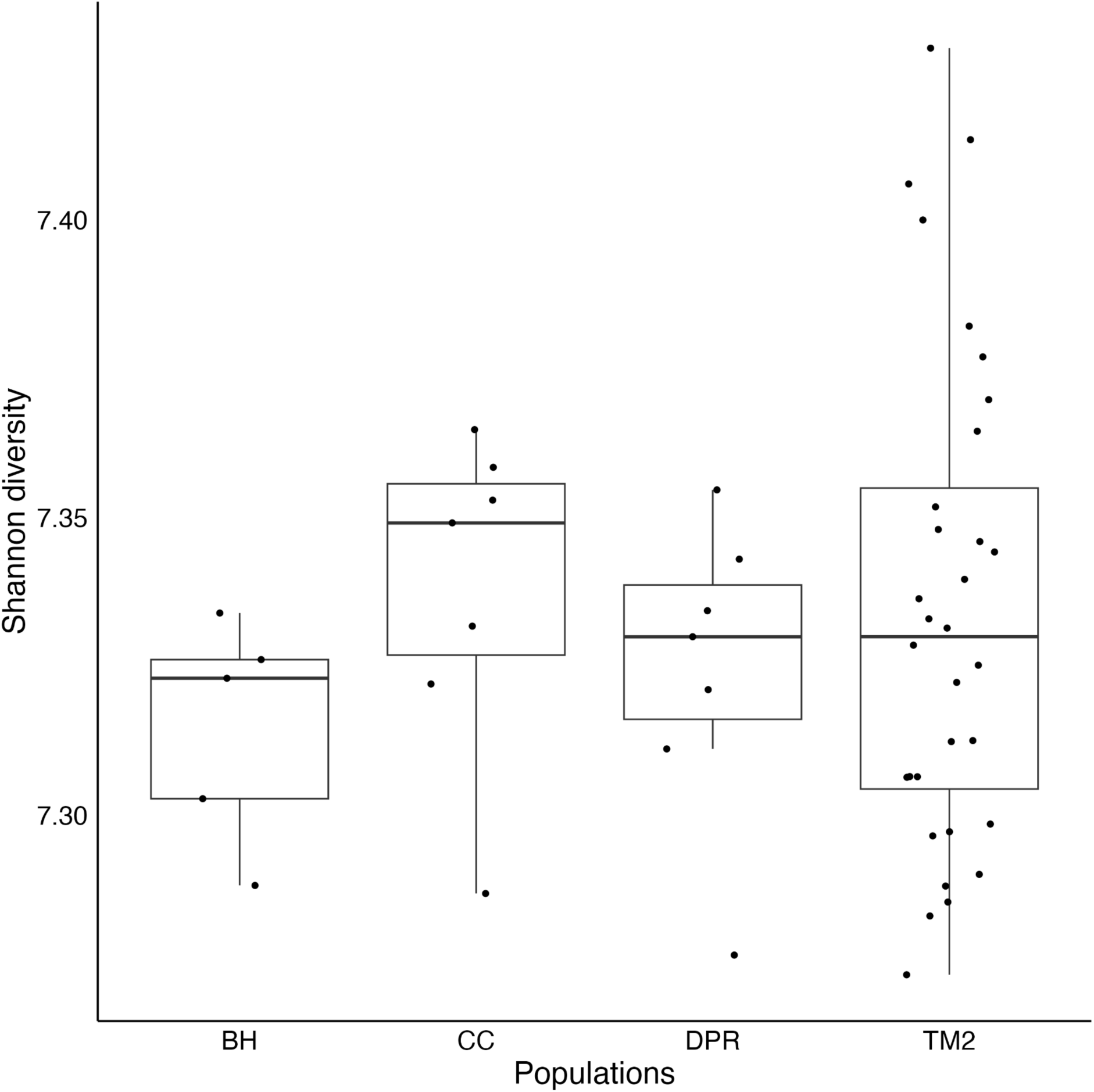
Shannon diversity index calculated on the reflectance data for all the samples used in the study. Each dot represents an individual sample for each population.

**Fig. S4:**
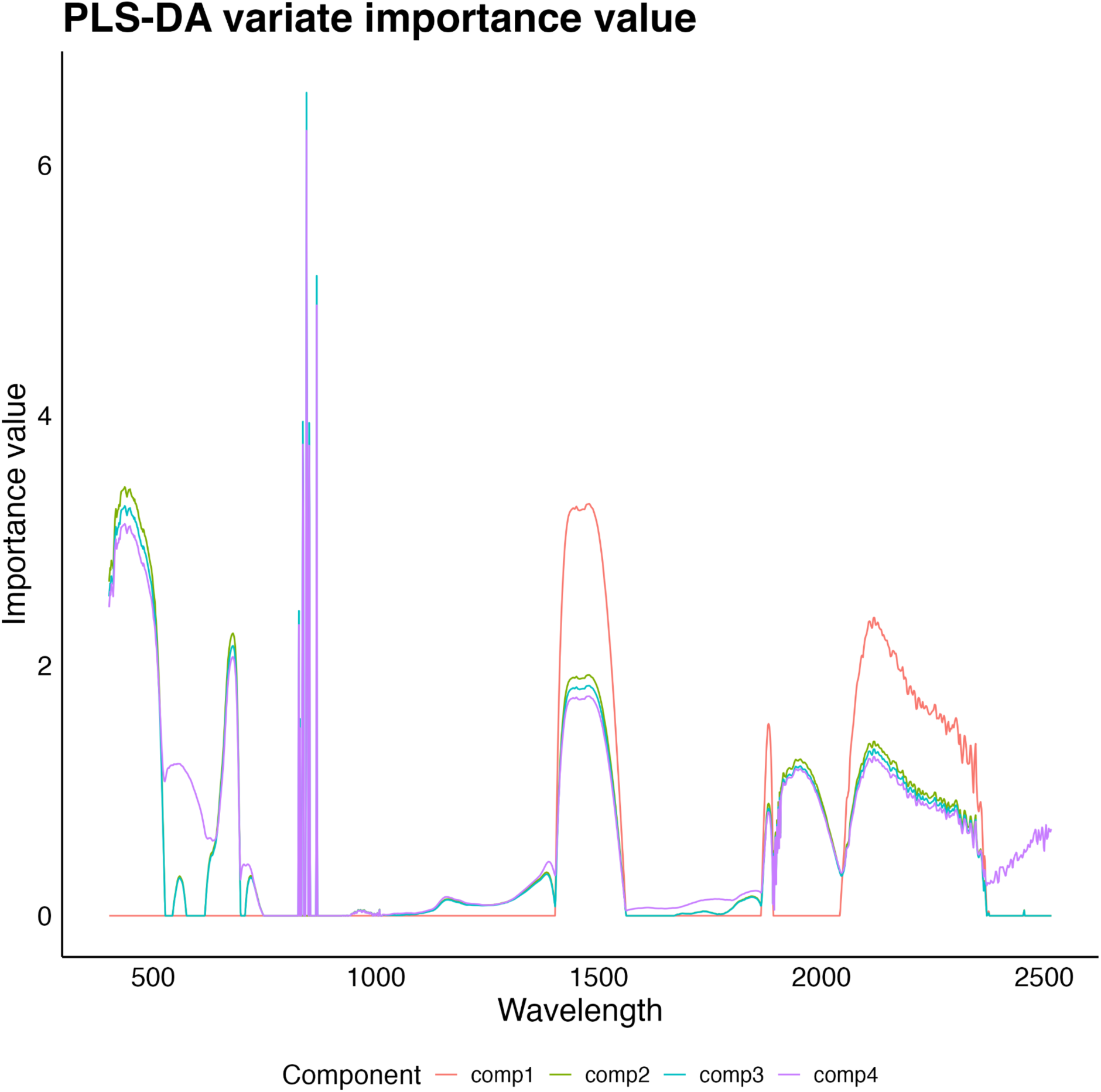
Importance value of each wavelength across the 4 PLS-DA variates from Fig. 1A.

## References

Albert R, Barabási A-L. 2002. Statistical mechanics of complex networks. Reviews of Modern Physics 74: 47–97.

Anderson JT, DeMarche ML, Denney DA, Breckheimer I, Santangelo J, Wadgymar SM. 2025. Adaptation and gene flow are insufficient to rescue a montane plant under climate change. Science.

Anderson JT, Gezon ZJ. 2015. Plasticity in functional traits in the context of climate change: a case study of the subalpine forb Boechera stricta (Brassicaceae). Global Change Biology 21: 1689–1703.

Anderson JT, Inouye DW, McKinney AM, Colautti RI, Mitchell-Olds T. 2012. Phenotypic plasticity and adaptive evolution contribute to advancing flowering phenology in response to climate change. Proceedings of the Royal Society B: Biological Sciences 279: 3843–3852.

Armbruster WS, Pélabon C, Bolstad GH, Hansen TF. 2014. Integrated phenotypes: understanding trait covariation in plants and animals. Philosophical Transactions of the Royal Society B: Biological Sciences 369: 20130245.

Asner GP, Martin RE, Anderson CB, Knapp DE. 2015. Quantifying forest canopy traits: Imaging spectroscopy versus field survey. Remote Sensing of Environment 158: 15–27.

Bannari A, Morin D, Bonn F, Huete AR. 1995. A review of vegetation indices. Remote Sensing Reviews 13: 95–120.

Barabási A-L, Oltvai ZN. 2004. Network biology: understanding the cell’s functional organization. Nature Reviews Genetics 5: 101–113.

Bascompte J, Jordano P. 2007. Plant-Animal Mutualistic Networks: The Architecture of Biodiversity. Annual Review of Ecology, Evolution, and Systematics 38: 567–593.

Bontrager M, Worthy SJ, Cacho NI, Leventhal L, Maloof JN, Gremer JR, Schmitt J, Strauss SY. 2025. Herbarium specimens reveal a constrained seasonal climate niche despite diverged annual climates across a wildflower clade. Proceedings of the National Academy of Sciences 122: e2503670122.

Burnett AC, Anderson J, Davidson KJ, Ely KS, Lamour J, Li Q, Morrison BD, Yang D, Rogers A, Serbin SP. 2021. A best-practice guide to predicting plant traits from leaf-level hyperspectral data using partial least squares regression. Journal of Experimental Botany 72: 6175–6189.

Caplan S, Huemmrich KF. 2025. Unveiling PACE OCI’s hyperspectral terrestrial data products. Remote Sensing Letters 16: 422–433.

Cavender-Bares J, Meireles JE, Couture JJ, Kaproth MA, Kingdon CC, Singh A, Serbin SP, Center A, Zuniga E, Pilz G, et al. 2016. Associations of Leaf Spectra with Genetic and Phylogenetic Variation in Oaks: Prospects for Remote Detection of Biodiversity. Remote Sensing 8: 221.

Cavender-Bares J, Schneider FD, Santos MJ, Armstrong A, Carnaval A, Dahlin KM, Fatoyinbo L, Hurtt GC, Schimel D, Townsend PA, et al. 2022. Integrating remote sensing with ecology and evolution to advance biodiversity conservation. Nature Ecology & Evolution 6: 506–519.

Cawse-Nicholson K, Chadwick KD, Brodrick PG, Kiper M, Thompson DR, Schimel D, Miller CE, Townsend PA, Alves LF, Shiklomanov AN, et al. 2025. Intrinsic dimensionality as a metric for temporal plant diversity evaluation: Case study from the SHIFT campaign. Ecosphere 16: e70213.

Chen Y, Monks L, Rubio VE, Cox AJ, Swenson NG. 2025. Linking leaf hyperspectral reflectance to gene expression. Communications Earth & Environment 6: 694.

Cheverud JM. 1996. Developmental Integration and the Evolution of Pleiotropy. American Zoologist 36: 44–50.

Davidson EH, Erwin DH. 2006. Gene Regulatory Networks and the Evolution of Animal Body Plans. Science 311: 796–800.

Des Roches S, Post DM, Turley NE, Bailey JK, Hendry AP, Kinnison MT, Schweitzer JA, Palkovacs EP. 2018. The ecological importance of intraspecific variation. Nature Ecology & Evolution 2: 57–64.

Edelsparre AH, Fitzpatrick MJ, Saastamoinen M, Teplitsky C. 2024. Evolutionary adaptation to climate change. Evolution Letters 8: 1–7.

Espinosa-Soto C, Wagner A. 2010. Specialization Can Drive the Evolution of Modularity. PLOS Computational Biology 6: e1000719.

Féret J-B, Berger K, de Boissieu F, Malenovský Z. 2021. PROSPECT-PRO for estimating content of nitrogen-containing leaf proteins and other carbon-based constituents. Remote Sensing of Environment 252: 112173.

Féret J-B, Boissieu F de. 2024. prospect: an R package to link leaf optical properties with their chemical and structural properties with the leaf model PROSPECT. Journal of Open Source Software 9: 6027.

Flint LE, Flint AL, Stern MA. 2021. The basin characterization model—A regional water balance software package. U.S. Geological Survey.

Franks SJ, Weber JJ, Aitken SN. 2014. Evolutionary and plastic responses to climate change in terrestrial plant populations. Evolutionary Applications 7: 123–139.

Furbank RT, Tester M. 2011. Phenomics – technologies to relieve the phenotyping bottleneck. Trends in Plant Science 16: 635–644.

Gamon JA, Somers B, Malenovský Z, Middleton EM, Rascher U, Schaepman ME. 2019. Assessing Vegetation Function with Imaging Spectroscopy. Surveys in Geophysics 40: 489–513.

Gamon JA, Wang R, Gholizadeh H, Zutta B, Townsend PA, Cavender-Bares J. 2020. Consideration of Scale in Remote Sensing of Biodiversity. In: Cavender-Bares J, Gamon JA, Townsend PA, eds. Remote Sensing of Plant Biodiversity. Cham: Springer International Publishing, 425–447.

Gienapp P, Reed TE, Visser ME. 2014. Why climate change will invariably alter selection pressures on phenology. Proceedings of the Royal Society B: Biological Sciences 281: 20141611.

Gladstone-Gallagher RV, Hewitt JE, Siwicka E, Gammal JM, Brustolin MC, Norkko A, Pilditch CA, Thrush SF. 2023. Ecological network analysis of traits reveals variable response capacity to stress. Proceedings of the Royal Society B: Biological Sciences 290: 20230403.

Gremer JR, Chiono A, Suglia E, Bontrager M, Okafor L, Schmitt J. 2020a. Variation in the seasonal germination niche across an elevational gradient: the role of germination cueing in current and future climates. American Journal of Botany 107: 350–363.

Gremer JR, Wilcox CJ, Chiono A, Suglia E, Schmitt J. 2020b. Germination timing and chilling exposure create contingency in life history and influence fitness in the native wildflower Streptanthus tortuosus. Journal of Ecology 108: 239–255.

Hansen TF, Houle D. 2008. Measuring and comparing evolvability and constraint in multivariate characters. Journal of Evolutionary Biology 21: 1201–1219.

He N, Li Y, Liu C, Xu L, Li M, Zhang J, He J, Tang Z, Han X, Ye Q, et al. 2020. Plant Trait Networks: Improved Resolution of the Dimensionality of Adaptation. Trends in Ecology & Evolution 35: 908–918.

Hein NT, Ciampitti IA, Jagadish SVK. 2021. Bottlenecks and opportunities in field-based high-throughput phenotyping for heat and drought stress. Journal of Experimental Botany 72: 5102–5116.

Hereford J. 2009. A Quantitative Survey of Local Adaptation and Fitness Trade-Offs. The American Naturalist 173: 579–588.

Hoffmann AA, Sgrò CM. 2011. Climate change and evolutionary adaptation. Nature 470: 479–485.

Houle D, Govindaraju DR, Omholt S. 2010. Phenomics: the next challenge. Nature Reviews Genetics 11: 855–866.

Jacquemoud S, Baret F. 1990. PROSPECT: A model of leaf optical properties spectra. Remote Sensing of Environment 34: 75–91.

Jeong H, Tombor B, Albert R, Oltvai ZN, Barabási A-L. 2000. The large-scale organization of metabolic networks. Nature 407: 651–654.

Karabourniotis G, Liakopoulos G, Bresta P, Nikolopoulos D. 2021. The Optical Properties of Leaf Structural Elements and Their Contribution to Photosynthetic Performance and Photoprotection. Plants 10: 1455.

Kaur S, Kakani VG, Carver B, Jarquin D, Singh A. 2024. Hyperspectral imaging combined with machine learning for high-throughput phenotyping in winter wheat. The Plant Phenome Journal 7: e20111.

Klingenberg CP. 2008. Morphological Integration and Developmental Modularity. Annual Review of Ecology, Evolution, and Systematics 39: 115–132.

Kokaly RF, Asner GP, Ollinger SV, Martin ME, Wessman CA. 2009. Characterizing canopy biochemistry from imaging spectroscopy and its application to ecosystem studies. Remote Sensing of Environment 113: S78–S91.

Leimu R, Fischer M. 2008. A Meta-Analysis of Local Adaptation in Plants. PLOS ONE 3: e4010.

Li C, Czyż EA, Halitschke R, Baldwin IT, Schaepman ME, Schuman MC. 2023. Evaluating potential of leaf reflectance spectra to monitor plant genetic variation. Plant Methods 19: 108.

Li Y, He N. 2024. Innovations and prospectives of multidimensional trait integration. New Phytologist 244: 337–340.

Magney TS. 2025. Hyperspectral reflectance integrates key traits for predicting leaf metabolism. New Phytologist 246: 383–385.

Meireles JE, Cavender-Bares J, Townsend PA, Ustin S, Gamon JA, Schweiger AK, Schaepman ME, Asner GP, Martin RE, Singh A, et al. 2020. Leaf reflectance spectra capture the evolutionary history of seed plants. New Phytologist 228: 485–493.

Merilä J, Hendry AP. 2014. Climate change, adaptation, and phenotypic plasticity: the problem and the evidence. Evolutionary Applications 7: 1–14.

Messier J, McGill BJ, Enquist BJ, Lechowicz MJ. 2017. Trait variation and integration across scales: is the leaf economic spectrum present at local scales? Ecography 40: 685–697.

Murren CJ. 2012. The Integrated Phenotype. Integrative and Comparative Biology 52: 64–76.

Nicotra AB, Atkin OK, Bonser SP, Davidson AM, Finnegan EJ, Mathesius U, Poot P, Purugganan MD, Richards CL, Valladares F, et al. 2010. Plant phenotypic plasticity in a changing climate. Trends in Plant Science 15: 684–692.

Niinemets Ü. 2001. Global-Scale Climatic Controls of Leaf Dry Mass Per Area, Density, and Thickness in Trees and Shrubs. Ecology 82: 453–469.

Oksanen J, Simpson GL, Blanchet FG, Kindt R, Legendre P, Minchin PR, O’Hara RB, Solymos P, Stevens MHH, Szoecs E, et al. 2025. vegan: Community Ecology Package.

Olesen JM, Bascompte J, Dupont YL, Jordano P. 2007. The modularity of pollination networks. Proceedings of the National Academy of Sciences 104: 19891–19896.

Pau S, Slapikas R, Ho C-L, Bayliss SLJ, Donnelly RC, Abdullahi A, Helliker BR, Nippert JB, Riley WJ, Still CJ, et al. 2025. Hyperspectral leaf reflectance of grasses varies with evolutionary lineage more than with site. Ecosphere 16: e70257.

Pierrat ZA, Magney TS, Richardson WP, Runkle BRK, Diehl JL, Yang X, Woodgate W, Smith WK, Johnston MR, Ginting YRS, et al. 2025. Proximal remote sensing: an essential tool for bridging the gap between high-resolution ecosystem monitoring and global ecology. New Phytologist 246: 419–436.

Pieruschka R, Poorter H. 2012. Phenotyping plants: genes, phenes and machines. Functional plant biology: FPB 39: 813–820.

Preston RE. 1991. The Intrafloral Phenology of Streptanthus Tortuosus (brassicaceae). American Journal of Botany 78: 1044–1053.

Preston RE. 1994. Pollination Biology of Streptanthus Tortuosus (brassicaceae). Madroño 41: 138–147.

Proulx SR, Promislow DEL, Phillips PC. 2005. Network thinking in ecology and evolution. Trends in Ecology & Evolution 20: 345–353.

R Core Team. 2022. R: A Language and Environment for Statistical Computing. Vienna, Austria: R Foundation for Statistical Computing.

Rohart F, Gautier B, Singh A, Lê Cao K-A. 2017. mixOmics: An R package for ‘omics feature selection and multiple data integration. PLoS Computational Biology 13: e1005752.

Savolainen O, Lascoux M, Merilä J. 2013. Ecological genomics of local adaptation. Nature Reviews Genetics 14: 807–820.

Schlosser G, Wagner GP (Eds). 2004. Modularity in Development and Evolution. Chicago, IL: University of Chicago Press.

Schweiger AK, Cavender-Bares J, Townsend PA, Hobbie SE, Madritch MD, Wang R, Tilman D, Gamon JA. 2018. Plant spectral diversity integrates functional and phylogenetic components of biodiversity and predicts ecosystem function. Nature Ecology & Evolution 2: 976–982.

Schwinning S, Lortie CJ, Esque TC, DeFalco LA. 2022. What common-garden experiments tell us about climate responses in plants. Journal of Ecology 110: 986–996.

Serbin SP, Singh A, McNeil BE, Kingdon CC, Townsend PA. 2014. Spectroscopic determination of leaf morphological and biochemical traits for northern temperate and boreal tree species. Ecological Applications 24: 1651–1669.

Singh A, Serbin SP, McNeil BE, Kingdon CC, Townsend PA. 2015. Imaging spectroscopy algorithms for mapping canopy foliar chemical and morphological traits and their uncertainties. Ecological Applications 25: 2180–2197.

Song P, Wang J, Guo X, Yang W, Zhao C. 2021. High-throughput phenotyping: Breaking through the bottleneck in future crop breeding. The Crop Journal 9: 633–645.

Stouffer DB, Bascompte J. 2011. Compartmentalization increases food-web persistence. Proceedings of the National Academy of Sciences 108: 3648–3652.

Tay JK, Narasimhan B, Hastie T. 2023. Elastic Net Regularization Paths for All Generalized Linear Models. Journal of Statistical Software 106: 1–31.

Thenkabail PS, Smith RB, De Pauw E. 2000. Hyperspectral Vegetation Indices and Their Relationships with Agricultural Crop Characteristics. Remote Sensing of Environment 71: 158–182.

Tigano A, Friesen VL. 2016. Genomics of local adaptation with gene flow. Molecular Ecology 25: 2144–2164.

Ustin SL, Gamon JA. 2010. Remote sensing of plant functional types. New Phytologist 186: 795–816.

Wagner GP, Altenberg L. 1996. Perspective: Complex Adaptations and the Evolution of Evolvability. Evolution 50: 967–976.

Wagner GP, Pavlicev M, Cheverud JM. 2007. The road to modularity. Nature Reviews Genetics 8: 921–931.

Wang R, Gamon JA, Cavender-Bares J, Townsend PA, Zygielbaum AI. 2018. The spatial sensitivity of the spectral diversity–biodiversity relationship: an experimental test in a prairie grassland. Ecological Applications 28: 541–556.

Wang X, Ji M, Zhang Y, Zhang L, Akram MA, Dong L, Hu W, Xiong J, Sun Y, Li H, et al. 2023. Plant trait networks reveal adaptation strategies in the drylands of China. BMC Plant Biology 23: 266.

Wong CYS. 2023. Plant optics: underlying mechanisms in remotely sensed signals for phenotyping applications. AoB PLANTS 15: plad039.

Wong CY, Gilbert ME, Pierce MA, Parker TA, Palkovic A, Gepts P, Magney TS, Buckley TN. 2023. Hyperspectral Remote Sensing for Phenotyping the Physiological Drought Response of Common and Tepary Bean. Plant Phenomics 5: 0021.

Wright IJ, Reich PB, Westoby M, Ackerly DD, Baruch Z, Bongers F, Cavender-Bares J, Chapin T, Cornelissen JHC, Diemer M, et al. 2004. The worldwide leaf economics spectrum. Nature 428: 821–827.

